# Nova: A library for rapid development of mass spectrometry software applications

**DOI:** 10.1101/2025.05.06.652494

**Authors:** Michael R. Hoopmann, Christopher D. McGann, Christopher M. Rose, Devin K. Schweppe

## Abstract

Nova is a software library for the reading, writing, and management of mass spectrometry data natively in the C# language. It complements similar software libraries ubiquitously used in application development for C++, Python, and Java. Few software libraries and resources for mass spectrometry data analysis have been developed for C# relative to other languages. C#, however, remains the dominant language of choice for development of real time mass spectrometry (RTMS) analysis and instrument control applications, illustrating the need for native C# libraries when developing RTMS software. Nova provides fast, easy-to-use structures and classes built upon interfaces that support open community standards, and are easily extensible for current or future mass spectrometry software development needs.

## Introduction

Software applications are built upon a foundation of reliable software libraries that provide basic functions, such as input and output, data storage and manipulation, and user interfaces. Software libraries are modular, designed to be inserted into applications when necessary, and speed application development by eliminating the redundant creation of the same functions for every application that needs them. Among mass spectrometry software for proteomics, there are a wealth of open source and freely available libraries for application development in C++,^1–3^ Python,^4–11^ and Java.^12–14^ These libraries provide common functions such as reading mass spectrometry data files and performing spectral analysis and evaluation.

Use of C# as an application development language is perhaps less widespread than C++ or Python. However, C# is required for development of real time mass spectrometry (RTMS) applications that utilize the Instrument Application Programming Interface (IAPI) from Thermo Fisher Scientific.^15^ The IAPI enables real time spectral analysis and instrument control, which has been demonstrated through the creation of, for example, real time search and efficient quantitation applications.^16–19^ To access the IAPI, applications must be developed using the C# programming language. Although it is possible to integrate C++ and C# using the C++/CLI, as done with Skyline,^20^ the process is not trivial and not desirable to the software developer. It is also possible to access the IAPI with Python using a bridge to the C# IAPI.^21^ Alternatively, it is preferable to use native C# mass spectrometry software libraries to perform common processing required in mass spectrometry data analysis. CSMSL^22^ and mzLib^23^ are two examples of native C# libraries for opening and viewing spectral data and libraries. However, they represent the limited number of available tools for C# relative to other programming languages. Particularly for RTMS application development, which often requires efficient and dynamic software designs to keep pace with the speed of data acquisition, there is a need for fast and robust data structures and methods for spectral data analysis, designed natively for C#.

We have created a library of classes and interfaces, named Nova, to facilitate development of native C# applications for MS data analysis without the need for C++/CLI. Nova contains classes for reading, storing, inspecting, and writing mass spectrometry data. Additionally, the library contains lightweight utilities for interprocess communication between applications, opening the potential to establish dataflows between multiple software tools written in C#. Though developed with the goal of forming a strong foundation for RTMS application development, Nova is not specific to RTMS and can be used for general MS software applications. Establishing a native C# library for spectra management not only removes the need for redundant construction, but also provides a fast and efficient solution for building powerful MS analysis tools.

## Methods

Nova is divided into three packages: data management, input/output, and interprocess communication. Each package is managed within a structured and hierarchical namespace for easy access to the underlying object classes (Figure 1). Nova is open source and freely available at https://schweppelab.github.io/Nova, including documentation and code examples.

**Figure 1:**
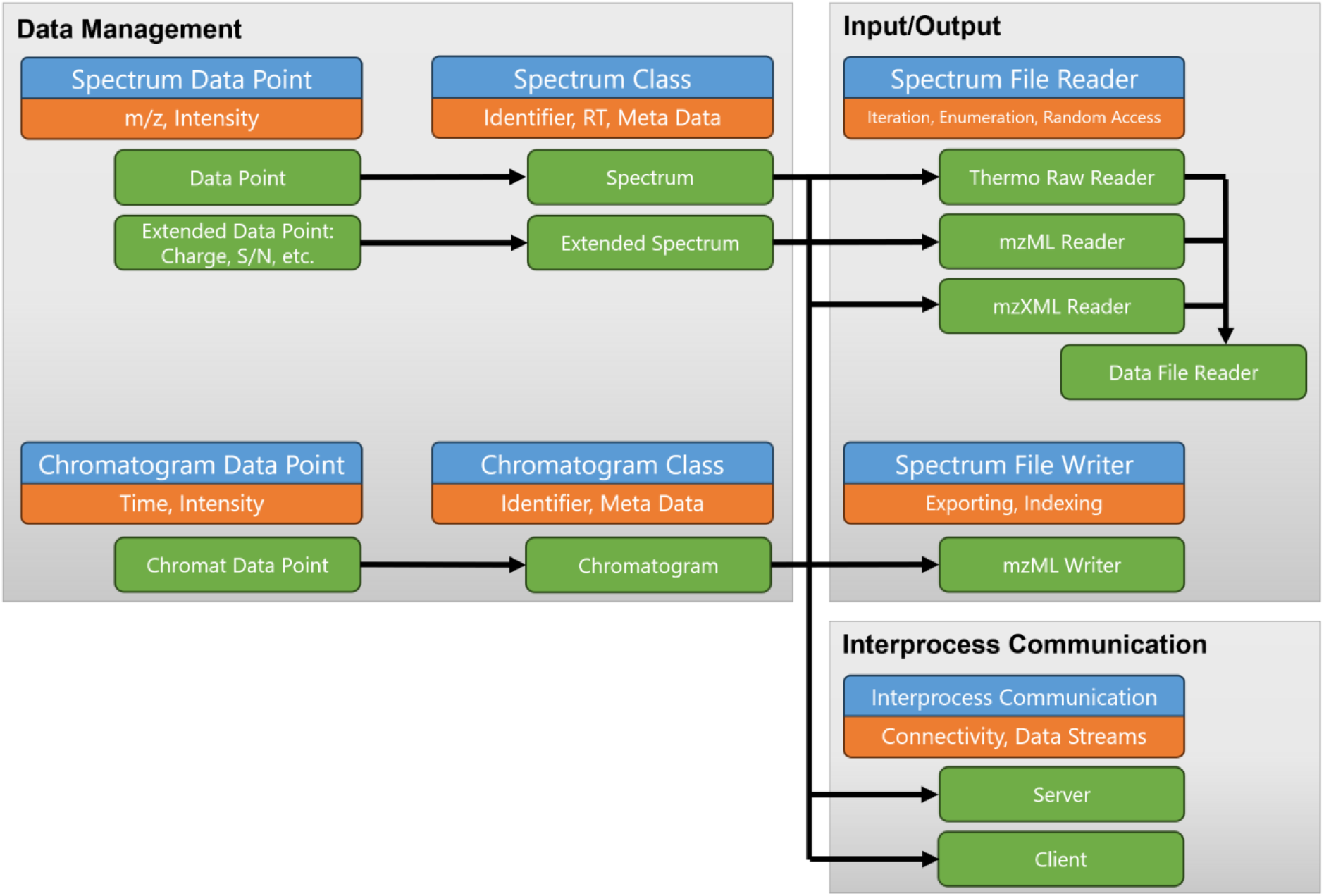
Nova architecture. Each of the three packages contains interfaces (blue) describing its contents and purpose (orange). From these interfaces, structures and classes (green) were developed to implement the functionality of Nova for mass spectrometry data analysis and interaction between the library packages.

### Data Management

The primary purpose of the data management package is to store relevant mass spectral information in data structures so that they may be managed and manipulated in software applications. Two principal class structures exist, *Spectrum* and *SpectrumEx* (extended spectrum), that are built upon a common foundation class containing relevant meta information and other properties pertaining to mass spectra. The *Spectrum* class contains a two-dimensional data array, generally for m/z and abundance values, but could also be used for other types of two-dimensional spectral such as chromatograms (time and abundance). These data are stored in arrays instead of lists typical of C# for fast access. The *SpectrumEx* class stores the same information as the *Spectrum* class, but also stores additional features for spectrum data points, such as baseline signal, noise threshold, peak resolution, and charge state. The intended use of this class is to manage and store processed spectra, for example, spectra converted to centroid peaks, each of which may have signal-to-noise ratio and precursor ion charge state assigned. Many downstream processing applications prefer processed spectra prior to use, and the *SpectrumEx* class provides a convenient means to store and manage processed spectra in larger analytical pipelines.

### Input/Output

The input/output (IO) package in Nova provides a means of reading and writing mass spectral data from and to a storage device. It works in conjunction with the data management package and uses the *Spectrum* and *SpectrumEx* object classes as its input and output. Mass spectra data are read iteratively, either sequentially or random-access, from a file reader object with a simple interface. A *Spectrum* object is returned, or if requested and supported by the data format, a *SpectrumEx* object containing the processed spectra. The reader currently supports open formats such as .mzML^24^ and .mzXML^25^, but can also read the vendor-native .raw format from Thermo Fisher Scientific using the RawFileReader library that gives access to centroid peak processing in conjunction with data reading. Nova also supports writing spectra back to file, to track and maintain any processing done to spectra during the course of analysis. The spectrum writer class stores spectra in the .mzML format. Though more robust tools exist that were designed explicitly for file conversion (e.g., ProteoWizard^2^), it is possible in Nova to quickly translate mass spectral data from one format to another by connecting the file input and output classes. Doing so with a processing method in between illustrates the ease with which data are loaded, transformed, and recorded, ready for use in any downstream analytical pipeline with Nova.

### Interprocess Communication

Interprocess communication occurs when data are shared between processes running on a computer. C# offers data pipes as a simple means of performing interprocess communication. Nova implements a named pipe wrapper, borrowing heavily from the established open source named-pipe-wrapper,^26^ to establish data traffic between applications. The easy-to-use design allows the user to define their own tokens to communicate the type of data, which is then transferred as a byte array. Methods on both the sending and receiving applications translate the byte arrays to and from class structures, using the tokens as guidance. Using the data pipes in Nova enables data to be transferred between applications in pipelines that use Nova without the need to read from or write to disk, as is the typical route in proteomics pipelines (e.g., reading .mzML data files). Particularly in the context of RTMS, which itself relies on interprocess communication between the instrument and user applications, it is possible to establish multiple applications into a pipeline capable of fast transfer of data between their processes to keep pace with the speed of data acquisition.

## Results & Discussion

Nova is a software library for fast reading, writing, and manipulation of mass spectrometry data using a C# development platform. Similar to libraries developed for other languages, Nova reads open, community developed mass spectrometry file formats, and also, in the case of Thermo Fisher Scientific data, native raw files. Spectral information is packaged into spectrum data structures that can be easily transferred between functions, or even between separate application processes. With barely a few lines of code, it is possible to read mass spectrometry data files and parse the collection of spectra within (Figure 2). These capabilities make Nova a solid foundation for further application development.

**Figure 2:**
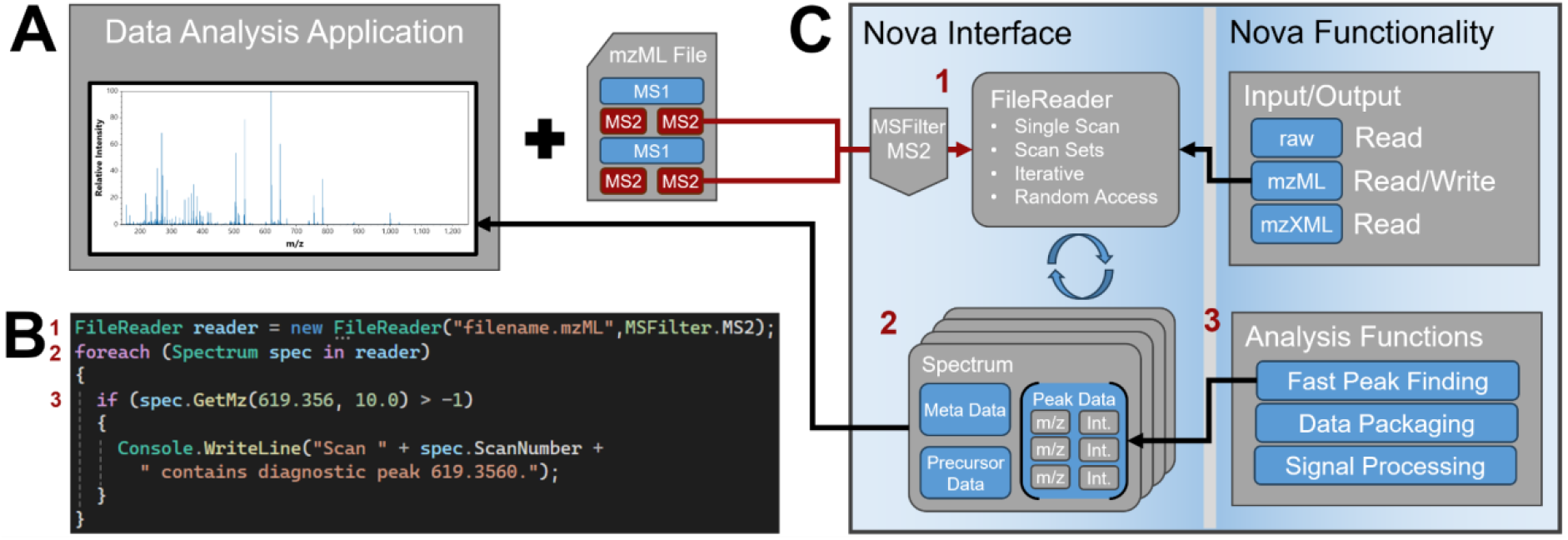
(A) Example of a simple application that processes mzML spectral data files. (B) Code to perform operations with Nova. Three lines of code that perform the application function are highlighted with red numbers. (C) Diagram of the Nova software library interface and functions, with inputs and outputs to the example application, and red numbers corresponding to the lines of code that perform the library operations. The first line of code (1) opens the file reader, which automatically identifies and accesses the correct Nova mzML file reader class, then sets a filter to only read MS/MS spectra from the input file. The second line of code (2) iterates each of those MS/MS spectra from the file, and the third line of code (3) uses the peak finding function to identify spectra containing a diagnostic peak m/z value.

**Figure 3:**
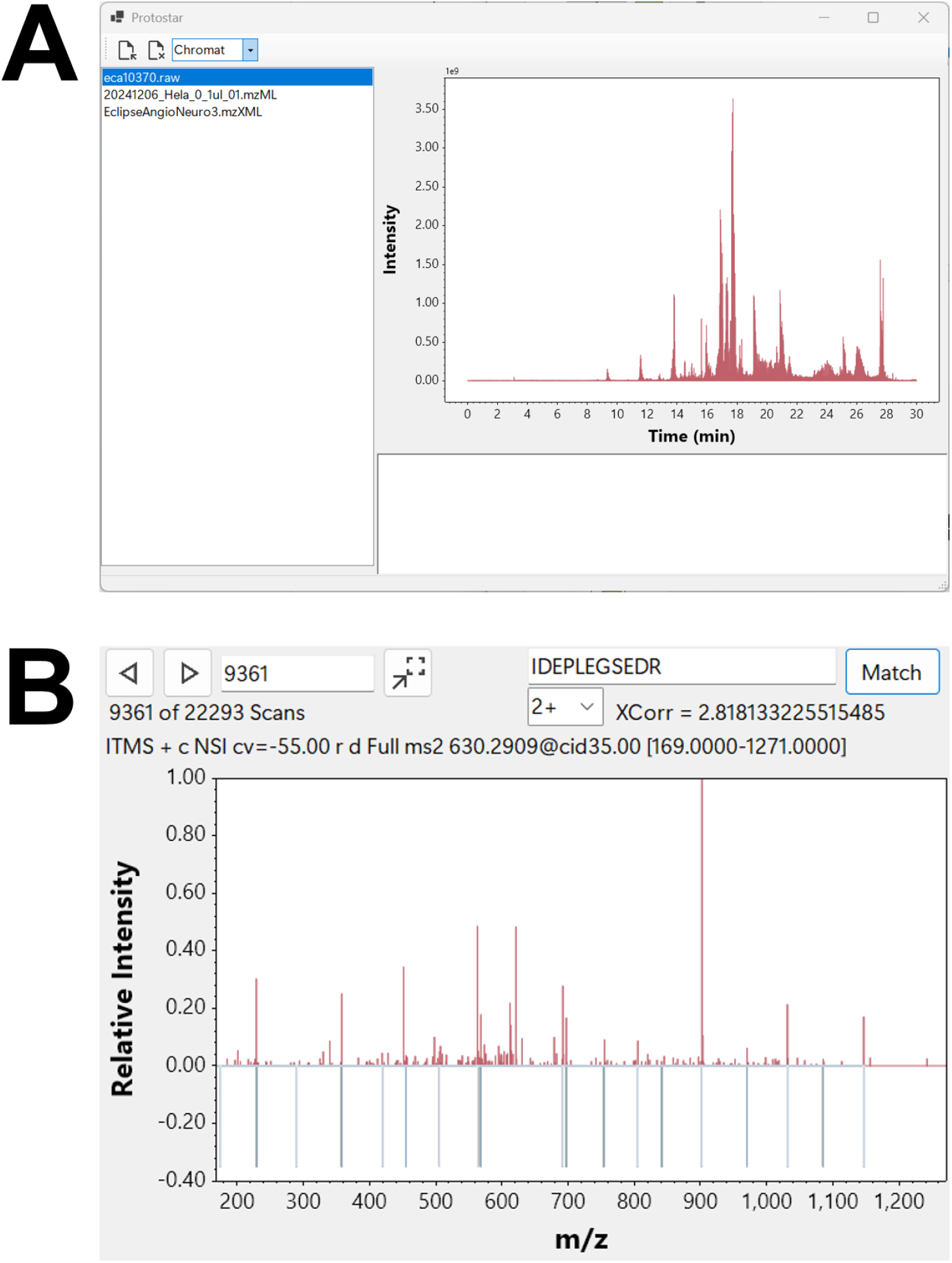
Protostar, a demonstration application using Nova. (A) The list of files being analyzed, with a chromatogram summary display. Multiple file formats are supported, including mzML, mzXML, and Thermo Fisher Scientific raw format. (B) Inspection of the data using the spectrum display. Peptide sequences supplied by the user produce an instantaneous PSM with Comet XCorr score value and corresponding butterfly plot of the fragment ion m/z values.

To exhibit the features of Nova, we created a simple demonstration application, Protostar, for visualizing mass spectrometry data. Using Protostar, it is possible to load multiple MS data files and contrast plots of their chromatography (Figure 1A). Files typically take a few seconds to load the first time, compounded slightly when loading the files simultaneously. Once loaded, the user can select and combine multiple total ion chromatograms from each file to inspect differences in chromatography, swapping between them instantaneously. Switching from chromatogram to spectrum mode provides deeper inspection of each spectrum within a selected file (Figure 1B). Navigating to any spectrum in the file can be done using the forward and backward buttons, or the jump-to button requesting any scan number within the data file. Spectra are loaded on-the-fly using the random-access data file reads that are built into Nova. Requesting any scan number requires only a single line of code. To demonstrate the ease of data management and manipulation, users can perform instant peptide spectral matches using the cross-correlation score (XCorr) from Comet.^27^ The underlying code transforms the spectrum structure to the fast cross-correlation structures and produces the score as would be observed during a spectral database search. Protostar returns the score, as well as a butterfly plot of the fragment ion m/z values for the requested peptide.

Nova contains two core aspects that maximize its utility for C# mass spectrometry software development, namely its spectrum-centric focus and its modularity. The basic data structures of Nova are its spectrum classes, mimicking the basic unit of each scan event in a mass spectrometer. All software that analyzes mass spectrometer data files must therefore process a spectrum scan event, and therefore maintaining this paradigm in the Nova software necessarily simplifies the design and speed with which analytical software can be written. It also provides a complement to other libraries developed for C# software development, such as mzLib^23^ and CSMSL^22^, which contain additional PSM, peptide, protein, FASTA, and spectral library class structures that define results and analytical features commonly found in mass spectrometry analytical pipelines. The modularity of Nova makes it easy to transform its spectrum objects to match the data structures of these libraries, to make use of their additional functionality, and vice versa, to extend the mzML file output functionality of Nova to the results of data processed with these libraries, for example. Though similar results can be obtained from other libraries, such as ProteoWizard,^2^ which offers a wide range of support for mass spectrometry data input and output using C++, it can pose a significant hurdle for novice developers in C#. Nova was designed to be lightweight, simple, and straightforward, for both novice and experienced developers using C#.

The architecture of Nova was deliberately designed to grow and adapt to the changing needs of mass spectrometry research. Interfaces describe common characteristics for each data structure and class, allowing interchange to occur in existing code that uses those interfaces, while enabling customization for future data structures and analyses that may be developed. Within those interfaces, unmanaged value types and structures are used where possible to minimize the boxing and unboxing of data to produce efficient performance, particularly when processing peak information, one of the heaviest computing tasks in mass spectrometry data analysis. Developers are encouraged to tailor Nova’s classes to their needs, using the existing interfaces and structures to ensure compatibility and performance.

## Conclusion

Nova is an open source, freely available software library for the management of mass spectrometry data and the development of analytical tools using the C# programming language. It provides a complement to similar software packages used when developing C++, Python, and Java applications for mass spectrometry data analysis. With the continued emergence of RTMS, which often requires C# software development, Nova supports native C# application development with the potential for use in RTMS studies.

